# Iron Metabolism and Adaptative Traits Associated with Virulence in *Enterobacter cloacae* Complex

**DOI:** 10.64898/2026.07.09.737523

**Authors:** Elizabeth W. Bugase, Tosin Y. Senbadejo, Lucas Ameng-Etego, Abiola Isawumi

## Abstract

Iron is an essential micronutrient that shapes host–pathogen interactions during infection. However, the contribution of iron to the virulence adaptation of the *Enterobacter cloacae* complex (ECC) remain poorly characterized. This study profiled the effects of iron on *E. roggenkampii* and *E. asburiae* clinical isolates. Growth kinetics were assessed in Luria–Bertani broth supplemented with varying iron concentrations and 5% sheep blood, and EDTA. Recovered strains were used for motility and antibiotic susceptibility assays. Phenotypic virulence trait of iron-naive and iron-recovered strains was determined using biofilm formation assays. Whole-genome sequencing was conducted to identify genetic determinants associated with iron acquisition and metabolism. Presence of iron increased bacterial growth, reduced antibiotic susceptibility, and enhanced biofilm formation. At higher iron concentrations, iron-recovered strains exhibited increased biofilm biomass, while there was a high biofilm formation with iron-naive strains at lower iron levels. Genomic analysis identified genes associated with ferrous and ferric iron transport, heme uptake, siderophore biosynthesis, and virulence-related functions, including adhesion and biofilm formation. These findings demonstrate that iron availability and prior exposure modulate ECC physiology and phenotypic traits associated with virulence, supporting a role for iron in shaping adaptive pathogenic potential.

**Graphical Abstract:** The influence of iron metabolism on virulence adaptation of *Enterobacter cloacae* complex

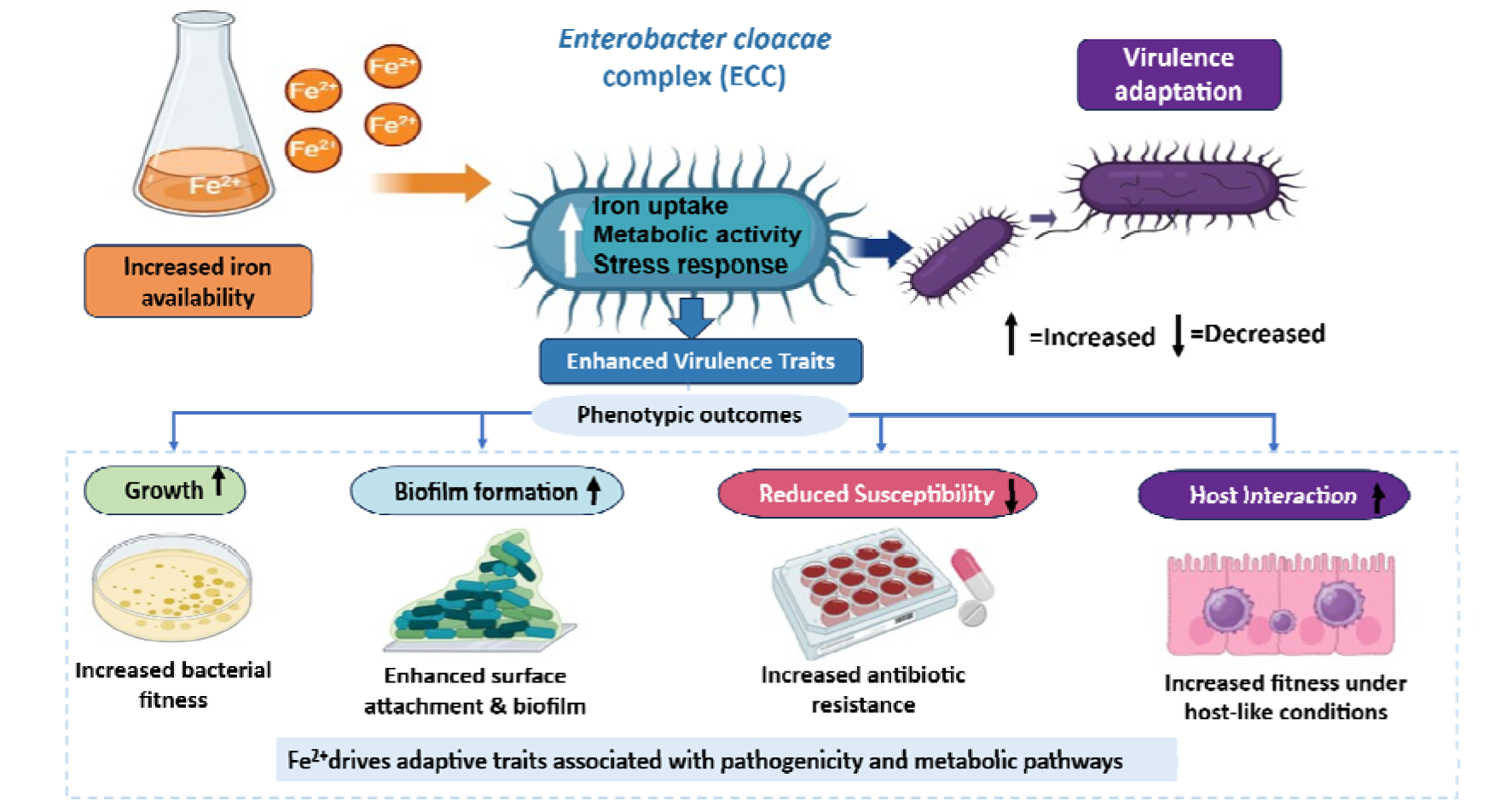

## Introduction

Iron is an important element involved in several biological processes including DNA synthesis, transport of oxygen and electrons for human cell survival (Abbaspour et al., 2014). Iron homeostasis in humans is tightly regulated because iron absorbed in the duodenum is utilized for heme synthesis in erythrocytes, while excess free iron can promote the generation of reactive oxygen species that may cause tissue damage if not efficiently controlled (Lieu et al., 2001; Winter et al., 2014).

In bacteria, iron is required for growth, replication, and other essential metabolic processes; Hence during bacterial infection, both invading bacteria and human host cells utilize iron for these vital cellular processes (Cassat & Skaar, 2013). Iron homeostasis is regulated by catalytic enzymes resulting in the generation of highly reactive oxygen species (Hussain et al., 2021). Through ‘nutritional immunity’, the host tightly regulates the amount of free iron available for utilization by invading bacteria (Hood & Skaar, 2012); helping the host to control the establishment of infections.

However, pathogenic bacteria such as the ESKAPE pathogens have developed mechanisms against host’s nutritional immunity to cause infections. This includes the use of siderophore machinery, heme factors, and heme-independent mechanisms (Hood & Skaar, 2012). The *Enterobacter cloacae* complex (ECC) is recognized as commensal organisms as well as opportunistic pathogens. The *Enterobacter* complex comprises a heterogenous group of closely related *Enterobacter* species, including *E. cloacae, E. asburiae, E. hormaechei, E. kobei, E. ludwigii E. roggenkampii, and E. bugandensis,* among others, with ongoing refinement of its taxonomic composition (Yaikhan et al., 2025). They are WHO priority pathogens, and as members of the ESKAPE group, implicated in nosocomial infections with high levels of antibiotic resistance (AMR)(Parau et al., 2019).

The ECC, like other commensals, requires iron for survival and adaptation and, as opportunistic pathogens, for host infection. It is possible that limitation of iron may trigger the expression of virulence-like factors involved in intravascular hemolysis, swarm motility, and biofilm formation, elevating the pathogenicity of ECC (Chu et al., 2010; Soria-Bustos et al., 2020; Zhang et al., 2016; Zhou et al., 2023). Previous studies have highlighted hemolytic activity in *E. cloacae* strains, particularly the production of a-hemolysin, which lyses erythrocytes ( Mezzatesta et al., 2012; Skals et al., 2009). Hemolysin is considered a putative virulence factor in ECC host infections and this can be exacerbated by co-occurrence with swarm motility (Simi et al., 2003).

Bacterial swarm motility is an important virulence-associated trait in ECC, contributing to bacterial migration, surface colonization, and host-pathogen interactions (Zegadło et al., 2023). During swarming, vegetative ECC cells differentiate into highly motile swarm cells that facilitate dissemination across surfaces and enhance colonization potential (Zegadło et al., 2023). In addition, ECC readily forms biofilms, a major virulence factor that enhances antimicrobial resistance (AMR), persistence, and survival under hostile environmental conditions (Warrier et al., 2021). Biofilm formation further contributes to the ability of ECC to cause chronic and nosocomial infections (Cai et al., 2024; Wutor et al., 2024). Despite increasing reports of multidrug-resistant (MDR) ECC, the physiological factors that modulate their virulence potential remain incompletely understood. In particular, the role of iron metabolism in shaping adaptive traits associated with virulence has not been fully elucidated. Given that iron availability is a critical environmental cue influencing bacterial physiology, understanding how ECC respond to varying iron conditions may provide insight into mechanisms underlying virulence expression and persistence.

Therefore, this study investigated the impact of different iron conditions (iron-rich, iron-limited, and host-mimicking environments) on adaptive traits associated with virulence in ECC isolated for ICUs of some selected hospitals in Ghana.

## Materials and Methods

### ECC strains and Growth

Archived ECC (i.e *Enterobacter roggenkampii* and *Enterobacter asburiae*) were retrieved from the ABISA bacterial culture library maintained in the, West African Centre for Cell Biology of Infectious Pathogens (WACCBIP), University of Ghana. Briefly, aliquots of frozen stock (-80 °C) were revived in sterile Luria Bertini (LB) broth and incubated (37°C 40 rpm) for 1 h before culturing on LB agar and incubated under the same conditions for 18 h. Pure colonies were subcultured on MacConkey agar and incubated (37 °C, 18 h). The strains recovered were used for subsequent assays.

### Media preparation and Growth Conditions for Iron-Stress Assays

#### Preparation of Host-mimicking Media

Luria–Bertani (LB) supplementation with sheep blood, 5%(v/v) supplies host-derived iron sources and induces hemolytic and oxidative stress, simulating bloodstream infections scenarios. To evaluate the impact of host-derived iron on growth and virulence, LB was supplemented with sheep blood (1-5%), with 5% used to mimic physiologically relevant heme-bound iron encountered during infection. Citrated sheep blood was obtained from the Animal Experimentation Unit of the Noguchi Memorial Institute of Medical Research, University of Ghana. Growth was assessed prior to iron stress assays. Together with FeSO_4_ supplementation and EDTA-mediated iron chelation, these media modifications simulated host-like conditions to examine ECC growth and virulence-associated traits.

#### Preparation of Iron-rich Media

FeSO_4_ supplementation provides exogenous iron to assess the impact of varying iron levels on the upregulation of iron acquisition systems and potential virulence in ECC. LB broth was supplemented with ferrous sulfate heptahydrate [FeSO_4._7H_2_O] to final concentrations of 0.1, 0.2, and 0.5 mM. To confirm that supplemented iron remained in the ferrous (Fe^2+^) state following autoclaving, sterile media were tested using 1,10-phenanthroline prior to use in growth assays.

#### Preparation of Iron-limiting Media

EDTA chelates residual iron in LB to mimic iron-limited conditions, enabling evaluation of adaptive responses such as production of siderophores and triggering of stress response pathways. Iron limited conditions were generated by supplementing LB with 20 μM of EDTA to chelate residual iron (∼10-16 μM) derived from tryptic and yeast extract components. EDTA was prepared from a 100 mM EDTA stock and filter-sterilized (22 μm), a concentration sufficient to induce iron limitation without significantly inhibiting bacterial growth (Eng et al., 2020).

### Iron Stress Assay

Overnight bacterial cultures were inoculated in fresh sterile LB broth and standardized to an optical density (OD) 0.05-0.1 used to setup four stress conditions. 1:100 dilution of the standardized bacterial culture in (1) LB supplemented with (0.1, 0.2, 0.5 mmol/L) of FeSO_4,_ (2) LB supplemented with 20 μM EDTA, (3) LB supplemented with 5% sheep blood and (4) LB broth without additives used as control to assess the growth profile of the strains. Growth kinetics of the strains were assessed by culturing strains in these experimental setups at 37 °C with shaking (40 rpm) and OD determined at 600 nm at 1 h intervals using the SkanIt Software version 4.1 for Microplate Readers RE, version 4.1.0.43, which was used for all assays in this experiment. Also, aliquots of the cultures were taken at different time intervals for quantification of intracellular iron.

### Iron Quantification

A standard calibration curve was generated for intracellular iron concentrations at different time points. Briefly, FeSO_4_ was prepared to obtain concentrations of 0.1, 0.2, 0.3, 0.5, 0.7, 0.9, and 1 mmol/L. Potassium acetate buffer (CH_3_COOK) and 1,10-Phenanthroline were added to Fe solutions with distilled H_2_O as a blank for the calibration. To quantify intracellular iron from the bacterial samples, aliquots of culture were centrifuged at 5000xg (10 min). Bacterial pellet was resuspended in 1 mL of buffer (CH_3_COOK), and 33.3 μL volume of HCl (4.1%) was added and incubated at room temperature (22 °C, 1 h) for complete lysis of bacterial cells. The pelleted cellular debris was harvested at 5000 x g (5 min), and 50 μL volume of sodium thiosulfate (Na_2_S_2_O_3_) was added to the supernatant containing intracellular Fe and incubated at 22 °C (20 min) to reduce ferric iron (Fe^3+^) to ferrous iron (Fe^2+^). After reduction, an equal volume of 1,10-Phenanthroline (1 mL) was added and the mixture was incubated at 22 [C (20 min) to form a ferroin [Fe (Phen)_3_]^2+^ complex with Fe^2+^ and absorbance measured at 510 nm.

### Hemolytic Activity

The hemolytic activity of the strains was determined qualitatively and quantitatively. For qualitative determination, briefly, sterile LB agar was supplemented with 5% citrated sheep blood. Pure colonies of ECC were streaked on the blood agar plates and incubated at 37 [C for 24 h. The formation of a clear zone/halo around colonies indicated complete lysis of erythrocytes (β-hemolysis), and partial hemolysis was indicated by a greenish or brownish zone (α-hemolysis). The quantitative analysis was carried out as previously described (Soria-Bustos *et al*., 2020) with minor modifications. Briefly, overnight cultures of ECC were diluted with fresh LB broth to an OD _600_ of 0.1 to standardize cell density for subsequent experiments. A 0.5 mL volume of the standardized culture was added to 0.5 mL of a 4% (v/v) RBC suspension. For the controls, 0.5 mL of 10% SDS was added to 0.5 mL of the RBC suspension as a positive control, while 0.5 mL of sterile PBS was used as a negative control. Following the 4 h incubation at 37 [C, the bacteria-RBC mix was centrifuged at 12000 x g for 1 min, and the hemoglobin released into the supernatant was quantified by measuring the absorbance at 450 nm. Following centrifugation, the bacterial-RBC pellet was used to prepare thin blood films, stained with Leishman’s stain, and examined microscopically for red blood cell morphological changes and hemolysis. Percentage hemolysis was calculated relative to the 10% SDS positive control.

### Minimum Inhibitory Concentration (MIC)

Micro broth dilution assay was used to determine the influence of iron uptake and utilization in resistance of ECC to meropenem and polymyxin B (PMB). Pure overnight cultures of iron stress-recovered ECC were standardized to 0.5 MacFarland (1 × 10[ CFU/mL,). Equal volumes (100 μL) of the standardized bacterial suspension and antibiotic solution prepared in two-fold serial dilutions of PMB (4-1024ug/ml) or Meropenem(5-1280ug/ml) were added to sterile 96-well microtiter plates and incubated at 37 °C for 24 h, and OD measured at 600 nm for MIC determination. MIC_90_ as used in this study is defined as the lowest antibiotic concentration required to inhibit ≥ 90% of bacterial population studied (Jones, 2002).

### Bacterial Surface Motility and Biofilm assays

Swarming motility assay was performed as described by Abban et al (2025) with slight modifications. Briefly, nutrient agar was prepared to a concentration of 0.7% (w/v). Sterilized media was cooled at 60 °C and transferred to Petri dishes (15 mL per plate). Overnight ECC cultures were standardized to OD_600_ of 0.2 to ensure a consistent inoculum. 5 μL of cultures was spotted in the middle of the agar plate and allowed to dry completely for 30 min before incubating at 37 [C for 24 h. Motility was assessed by measuring the diameter (mm) of the widest point of spread. For ECC recovered from iron utilization assays, strains were purified on MacConkey agar and cultured in LB broth to OD_600_ of 0.2 before spotting swarm media plates with 5 μL of standard culture for smarming assay.

Biofilm formation was assessed using the 96-well microtiter plate. A 200 μL volume of overnight cultures normalized to OD_600_ 0.1 in LB broth was transferred to 96-well plates and incubated at 37 °C, 48 h. Following incubation, plates were washed using sterile distilled water to remove spent media and loosely adherent bacteria. Plates were air-dried for 30 min, stained with 0.1% (w/v) crystal violet solution, and incubated at 22 C for 30 min. Plates were washed with sterile distilled water, air-dried and quantitatively assessed with 200 μL of 96% ethanol and absorbance determined at 570 nm. ECC recovered from iron assays and were equally subjected to biofilm formation assays. Also, iron-naive ECC pure cultures were diluted with media containing different LB supplementations (0.1, 0.2, 0.5 mM of FeSO_4,_ 20 μM EDTA, 5% sheep blood, and LB broth only to serve as a control) to OD_600_ 0.1 and subjected to biofilm formation as described above. This assesses how acute exposure and prior sensitization to different iron conditions affect the biofilm forming abilities of ECC. Absorbances representing means from biological replicates were plotted using heatmaps to evaluate differences in adherent biofilm formation.

#### Whole Genome Sequencing and Bioinformatic Analysis

Genomic DNA was extracted from bacterial isolates using the Phenol-chloroform standard protocol and submitted for sequencing at the WACCBIP Next Generation Sequencing Laboratory. Illumina workflows were used to generate paired-end sequencing reads in FastQ format, which were quality checked using FastQC and assembled *de novo* using SPAdes. The resulting draft genome assemblies were used for downstream analyses. Genome annotation was done using both Bakta (https://bakta.computational.bio/submit) and RAST (https://rast.nmpdr.org/rast.cgi) to generate standardized reproducible and functional subsystem-based annotations. Annotated GenBank files by Bakta were used as primary output for downstream visualization.

Iron acquisition and metabolism-related genes were identified from the annotated genomes by manual curation of annotation outputs and functional categories associated with iron transport, siderophore biosynthesis and uptake, heme/hemin utilization, and iron-dependent metabolic processes. Contigs were screened for the number of iron-associated genes, and the contig containing the most iron acquisition and metabolism genes were selected for visualization using Proksee (https://proksee.ca/). Circular genome maps were generated to display coding sequences (CDSs), iron acquisition and metabolism genes, and predicted horizontally acquired regions identified using the AlienHunter module implemented in Proksee. Contig 2 and 14 of *E. asburiae* and *E. roggenkampii*, which harbored the highest number of iron-related genes, were selected for visualization; thus, the maps represent a single contig rather the complete genome assembly.

Antimicrobial resistance (AMR) genes were identified using Resistance Gene Identifier (RGI) module in the Comprehensive Antibiotic Resistance Database (CARD). Only ‘strict’ and ‘perfect’ hits using the criteria; sequence identity ≥ 95% and coverage ≥ 80%. These stringent criteria were applied to minimize false positive predictions.

Virulence factors were identified using the Virulence Factor Database (VFDB) through BLAST-based sequence similarity searches against curated virulence gene datasets. Hits were filtered using an E-value threshold of ≤ 1×10^-5^, sequence identity ≥ 70% and query coverage ≥ 50%. These relaxed thresholds were applied to improve detection of divergent virulence gene homologs in draft genome assemblies, where gene fragmentation and sequence variability may reduce detectability under more stringent cut-offs.

#### Data Analysis and Statistics

All experiments were conducted with two biological and technical replicates to ensure reproducibility. Data from experiments were analyzed using GraphPad Prism v9.0.121 and plots of means with standard error bars representing deviations from the mean used to show experimental variability. Statistical significance in motility of ECC post iron assays was assessed using two-way ANOVA with Dunnett’s multiple comparisons test.

Comparative analyses of iron acquisition systems, resistance determinants, and virulence-associated genes were performed using gene counts and functional categories from annotation and blast analyses. Data was summarized and displayed using comparative charts, circular genome visualizations to highlight both shared and strain-specific features.

## RESULTS

### Enhanced growth of ECC in 5% sheep, EDTA and increased FeSO_4_ media supplementation

ECC strains exhibited enhanced growth with increasing concentration of sheep blood, with 5% supplementation producing the highest growth (Fig.1 A & B). This is consistent with utilization of blood-derived nutrients, potentially including heme-associated iron. This observation is consistent with bacteria utilizing iron from erythrocytes, which are major reservoirs of iron during bloodstream infections. It suggests that exposure to host blood and blood-derived factors in vivo could support bacterial growth and may enhance pathogenic potential of ECC. The growth of *E. asburiae* (Fig.1.A) at 2% sheep blood showed an increased duration of lag phase compared to the 1%. This suggests a possible initial high fitness cost and a subsequent adaptation to the various components of blood through the utilization of blood-derived iron and other nutrients reflected by enhanced bacterial growth at 24 h. This informed the choice of 5% sheep blood supplementation for subsequent iron stress assays.

**Fig. 1.**
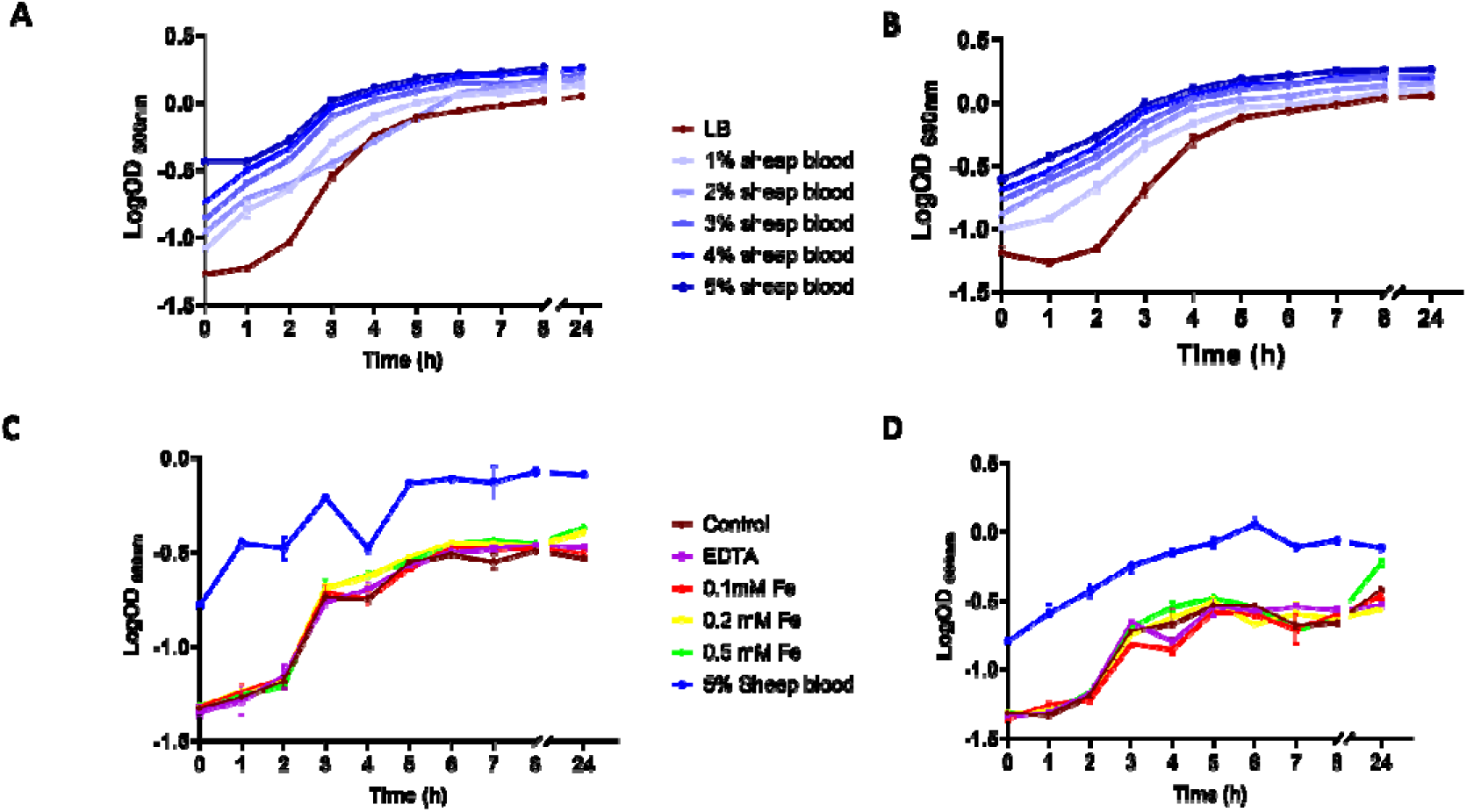
Growth profiles of *E. asburiae* and *E. roggenkampii* under different growth conditions.

The ECC strains exhibited enhanced growth in the media supplemented with 5% sheep blood (Fig. 1.C & D). FeSO_4_ (0.5 mM) supplemented media also support the growth of ECC strains as compared to 0.1-0.2 mM FeSO_4,_ EDTA and broth-only control. There is a general trend of enhanced bacterial growth with increasing Fe concentrations. This further supports the potential utilization of iron and blood-derived iron and nutrients for survival, highlighting the pathogenic potential of the ECC strains. Growth was not inhibited under iron-limiting conditions at 20 μM, indicating adaptation of ECC strains to low iron availability.

In panel A and B, LB broth was supplemented with 1-5% sheep blood showing increasing gradient of 24 h growth curves colour coded blue each with LB broth only control colour coded brown. In panels C and D growth curves showing similar curves in media supplemented with 0.1, 0.2 and 0.5 mM FeSO_4,_ 5% sheep blood and 20 μM EDTA. *(brown-LB broth only control, purple-20* μ*M EDTA, red-0.1 mM FeSO_4_, yellow-0.2 mM FeSO_4_, green-0.5mM FeSO_4_, blue-5% sheep blood). Error bars indicate the standard errors of the mean.* An axis break between the 8h and 24h measurements reflect extended incubation prior to the final measurement.

### ECC strains accumulate Fe²U intracellularly and display hemolytic potential

Iron supplementation resulted in time and concentration-dependent changes in intracellular iron levels (Fig. 2A & B). Under Fe²LJ-supplemented conditions (0.1, 0.2, & 0.5 mM FeSO_4_), intracellular iron increased during early time points, reaching peak levels at approximately 3 h, followed by a decline at later time points (6–24 h). The magnitude of this increase varied with iron concentration, with lower concentrations (0.1-0.2 mM) generally showing more pronounced transient accumulation compared to 0.5 mM. EDTA treatment reduced intracellular iron levels in both strains. However, the magnitude of reduction differed. In *E. asburiae* (Fig. 2A), intracellular iron remained consistently low across all time points under EDTA conditions. In contrast, *E. roggenkampii* (Fig. 2B) retained detectable intracellular iron despite EDTA treatment, indicating a comparatively reduced impact of iron chelation. In the 5% sheep blood supplementation, both strains exhibited elevated intracellular iron at early time points, followed by a marked decrease over time, approaching or falling below levels observed in Fe²LJ-supplemented conditions by 6 h (Fig. 2 A&B). While both strains displayed broadly similar uptake dynamics, differences were observed in the magnitude and rate of change in intracellular iron levels, indicating potential strain-specific variation in iron acquisition and regulation.

**Fig 2.**
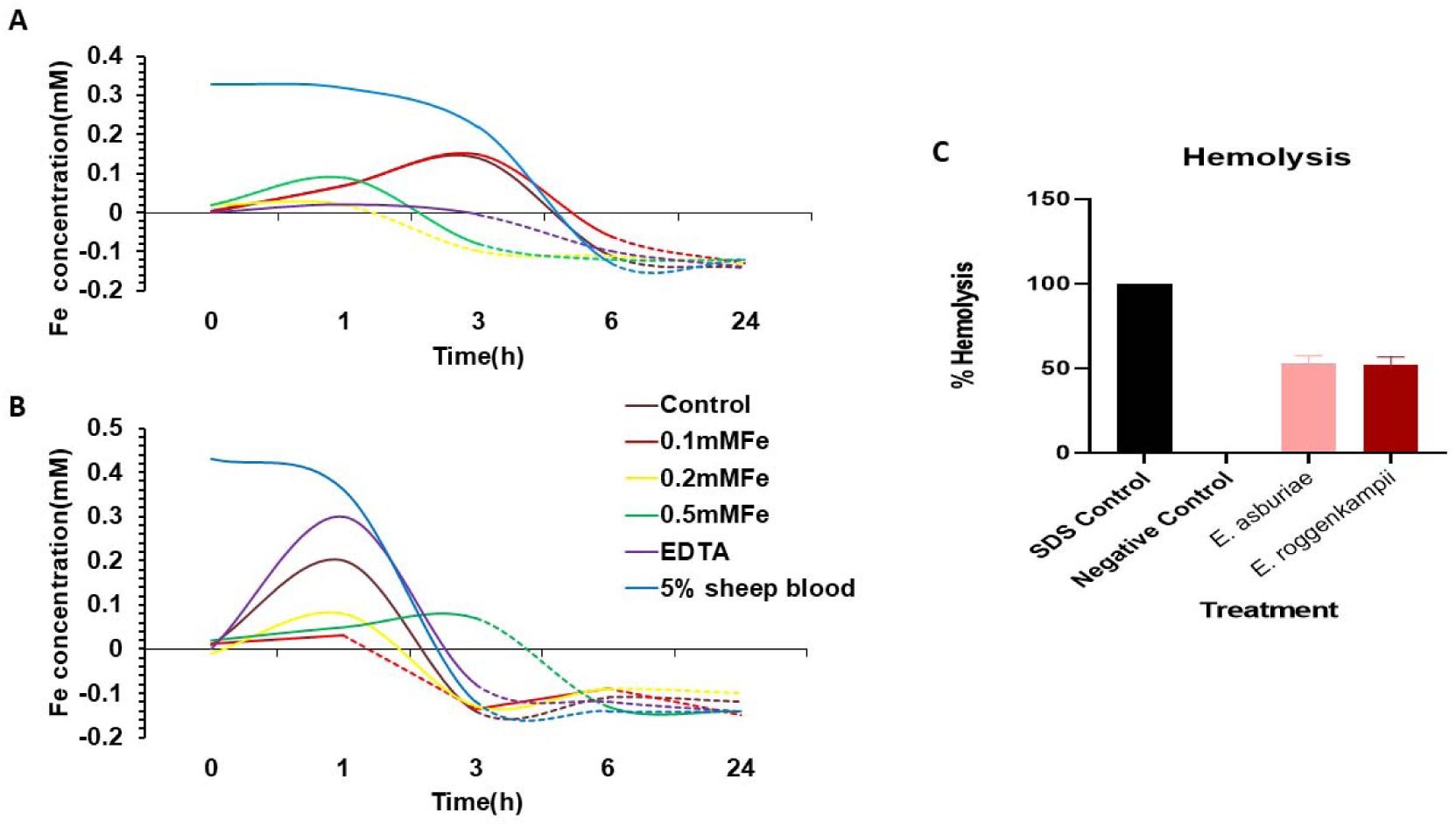
Intracellular iron concentrations in *E. asburiae* and *E. roggenkampii* post iron uptake under different iron concentrations. In panel A and B, LB was supplemented with 0.1,0.2 and 0.5mM FeSO_4,_ 5% sheep blood and 20 μM EDTA. *(brown-LB broth only control, purple-20* μ*M EDTA, red-0.1mM FeSO_4_, yellow-0.2mM FeSO_4_, green-0.5mM FeSO_4_, blue-5% sheep blood).* Panel C shows partial hemolysis by *E. asburiae* (pink) and *E. roggenkampii* (red) relative to 10% SDS positive control (black) and PBS negative control. *Error bars indicate the standard errors of the mean*.

ECC strains exhibited partial erythrocyte lysis, indicating their ability to interact with and damage erythrocytes and gain access to host-derived iron when cultured on chocolate agar. Microscopy of thin smears after incubation with RBCs confirmed partial hemolysis, indicated by cellular debris and remnants of bacterial cells when compared to PBS control, showing intact biconcave RBCs with areas of central pallor (supplementary). Fig. 2. Panel C indicates partial lysis of erythrocytes was observed by ECC strains as indicated by ≈50% hemolysis compared to the complete hemolysis (100%) in10% SDS positive control, and zero hemolysis in PBS negative control. This may suggest the ability of the ECC strains to utilize blood-derived iron through the lysis of erythrocytes.

### Iron utilization modulates Polymyxin B and Meropenem MICs in ECC strains

ECC strains demonstrated increased resistance to polymyxin B (PMB) post-iron uptake as indicated by high MIC_90_ (Fig. 3. A & B). The parental *E. asburiae* grown in polymyxin-LB broth (control) had a MIC_90_ of 16 μg/ml with ≤10% bacterial population by CFU/ml. *E. asburiae* strains grown in iron-supplemented media (0.1, 0.2, 0.5 mM FeSO_4,_ and 5% sheep blood) and iron-depleted media (EDTA) have MIC_90_ ≥1024 μg/ml (Fig. 3A). Similarly, *E. roggenkampii* parental control strain has a MIC_90_ of 16 μg/ml as compared to 8 μg/ml of EDTA and ≥1024 μg/ml for iron supplemented media (Fig. 3B). The influence of iron supplement and iron-depleted conditions on the change in susceptibility profiles of ECC strains to PMB suggests a possible role of iron metabolism in resistance. There was a general trend of an increase in bacterial population with increasing PMB concentration, preceded by an initial decrease in bacterial population. The observation of this non-monotonic response in contrast to a monotonic decline in response to increasing concentrations of PMB may be suggestive of an underlying resistance or survival mechanism in a subpopulation.

**Fig. 3.**
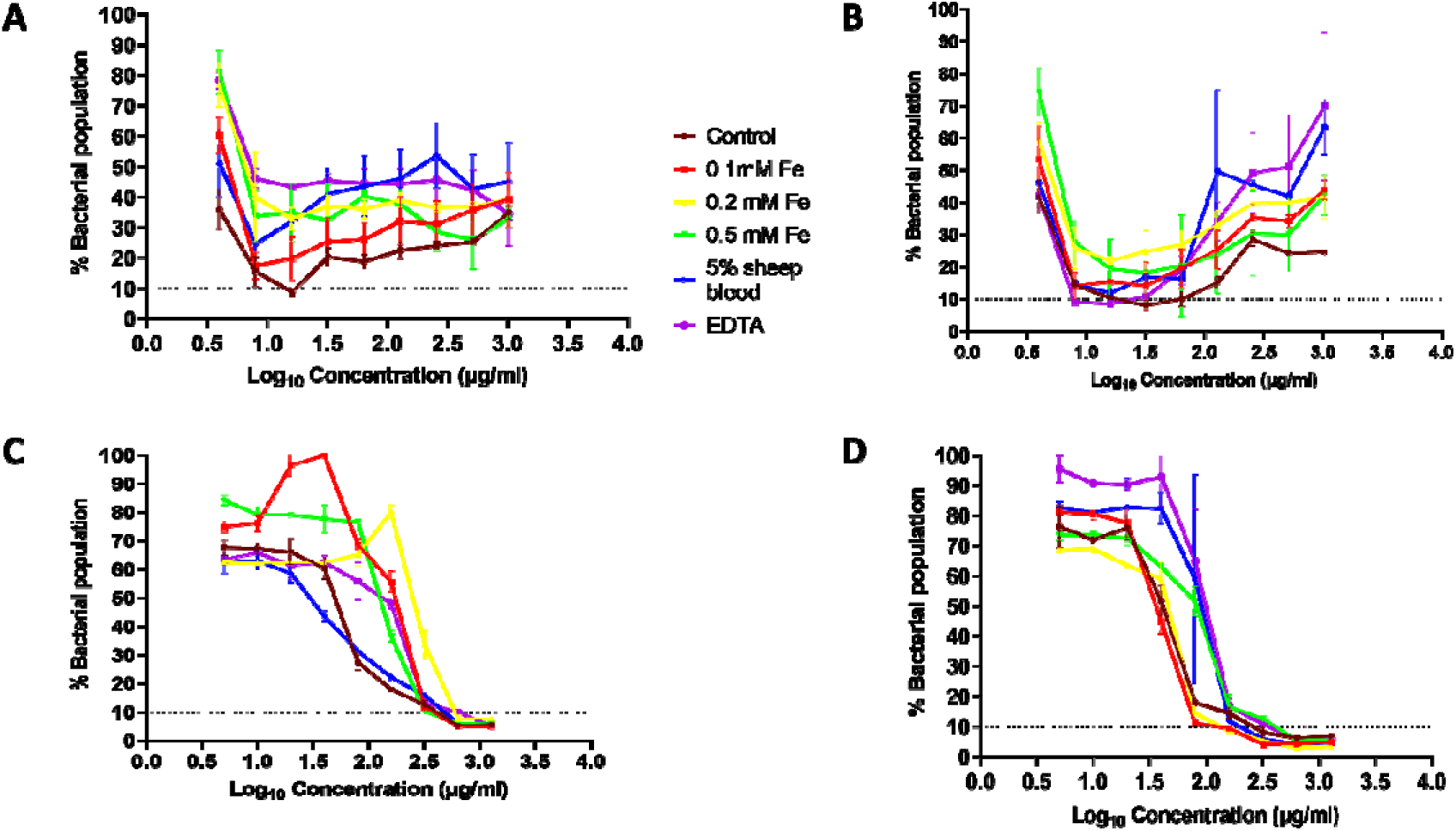
Minimum Inhibitory Concentrations of *E. asburiae* and *E. roggenkampii* against PMB and meropenem. In panel A *(E. asburiae*) and B (*E. roggenkampii*) response to standard and increasing concentrations of PMB post iron uptake. In panel C (*E. asburiae*) and D (*E. roggenkampii*) response to standard and increasing concentrations of meropenem post iron uptake. Concentrations are plotted on a log_10_ scale to reflect serial dilution series. *(brown-LB broth only control, purple-20* μ*M EDTA, red-0.1mM FeSO_4_, yellow-0.2mM FeSO_4_, green-0.5mM FeSO_4_, blue-5% sheep blood). Error bars indicate the standard errors of the mean*.

ECC displays high resistance to meropenem (Fig. 3. Panel C & D). *E. asburiae* strains in the various conditions had MIC_90_ of 640 μg/ml. However, the strains in 0.1, 0.2, 0.5 mM FeSO_4_ and 20 μM EDTA had higher bacterial population relative to the control. This suggests that iron-depleted and iron-rich conditions increase tolerance to higher meropenem concentrations. Also, *E. asburiae* in 5% sheep blood had an increased survival against meropenem indicated by a shift to the right which preceded an initial sensitive population compared to the control. Thus, suggesting a high fitness cost and later tolerance to other blood-derived components (Fig. 3C). *E. roggenkampii* preconditioned in 0.1 and 0.2 mM FeSO_4_ are relatively more sensitive to meropenem with a MIC_90_ of 160 μg/ml as compared to the broth only control 320 μg/ml). However, *E. roggenkampii* in 5% sheep blood supplement and 20 μM EDTA showed more tolerance to meropenem with a MIC_90_ of 320 μg/ml. The 0.5 FeSO_4_ pre-conditioning increased the strain resistance to meropenem with a MIC_90_ of 640 μg/ml (Fig. 3D). These findings highlight the influence of iron metabolism in adaptation of resistance; however, the data points to strain specific effect.

### Iron metabolism influences motility and biofilm formation in ECC strains

Motility plates were not supplemented with iron or chelator and inocula were not washed; thus, effects reflect pre-conditioning rather than on-plate iron availability. ECC strains are motile and demonstrated altered motility post iron uptake. *E. asburiae* recovered from 0.5 mM FeSO_4_ and 20 μM EDTA showed increased swarm motility compared to the control. However, these are not statistically significant (*p*LJ*0.05*) (Fig. 4A). *E. roggenkampii* grown in LB broth only (control) demonstrated increased swarm motility as compared to 0.1-0.2 mM FeSO_4_ and 5% sheep blood supplement. An increase in swarm motility was observed in the 0.5 mM FeSO_4_ and 20 μM EDTA conditions relative to the control and the difference was statistically significant (*p*≤*0.0001*) (Fig. 4 B). Pre-exposure to high iron or EDTA increased swarming in *E. roggenkampii* and variably in *E. asburiae*; iron availability during motility was not tested. Like *E. roggenkampii*, *E. asburiae* grown in LB broth only (control) demonstrated increased swarm motility compared to 0.1 mM FeSO_4_ and 5% sheep blood but decreased swarm motility to 0.2 mM FeSO_4_ supplement, but the difference was not statistically significant. These findings suggest strains specific adaptation to swarming motility which could also facilitate dissemination of bacteria and virulence adaptation.

**Fig. 4.**
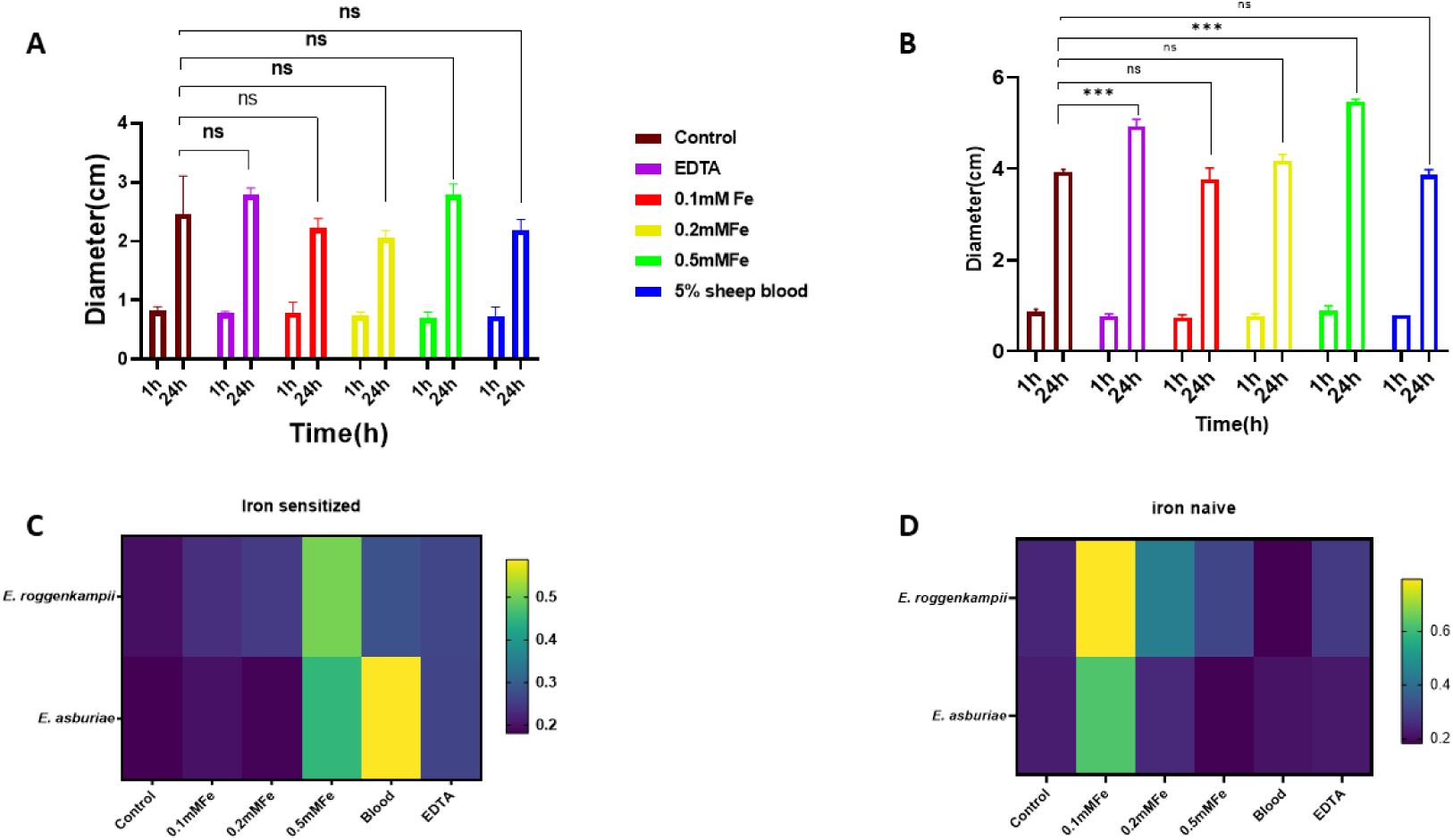
0. Swarming motility and biofilm formation in ECC strains. Diameter(cm) plotted to represent swarm motility at 1and 24h. *E. asburiae* (A) and *E. roggenkampii* (B) post iron uptake. *(brown-LB broth only control, purple-20* μ*M EDTA, red-0.1mM FeSO_4_, yellow-0.2mM FeSO_4_, green-0.5mM FeSO_4_, blue-5% sheep blood). Error bars indicate the standard errors of the mean.* Heat maps showing the biofilm biomasses of *E. asburiae* and *E. roggenkampii* with prior sensitization to different iron conditions (Fig. 4C) and biofilm formation without prior sensitization to same varying iron conditions (Fig. 4D). Gradient scale indicates Optical Density (OD) values at 570 nm post crystal violet standard biofilm formation. *Error bars indicate the standard errors of the mean*.

The formation of biofilm by ECC strains varies across different iron conditions, although crystal violet measures total stained biomass and may be influenced by growth/survival dynamics. Both iron-naive (without prior iron sensitization) and recovered strains were used in biofilm formation assays. Iron-recovered strains exhibited a dose-dependent increase in biofilm biomass, with the most robust biofilm formation occurring at the highest tested iron concentration. The total biomass of both *E. asburiae* and *E. roggenkampii* strains recovered from iron growth assays (iron sensitized) was increased in EDTA, and to a greater extent in 0.5 mM FeSO_4_ and 5% sheep blood supplement (Fig. 4C). The biofilm biomass of *E. asburiae* in 0.2 mM FeSO_4_ did not change but increased in *E. roggenkampii* sensitized in 0.2 mM FeSO_4._ Biofilm biomass of *E. roggenkampii* also increased in 0.1 mM FeSO_4_ as compared to the sensitized *E. asburiae* in 0.1 mM FeSO_4._ Biofilm formed by ECC with no prior iron exposed (iron naive) cells show a higher biomass in 0.1 mM FeSO_4_ supplementation compared to other conditions (Fig. 4 D). *E. roggenkampii* showed increased biofilm biomass in all conditions except in the 5% blood supplement which shows a decreased biofilm biomass compared to the control. *E. asburiae* in 0.5 mM FeSO_4_ showed decreased biofilm biomass compared to the 0.2 mM FeSO_4,_ 5% sheep blood and EDTA. Carryover of extracellular iron or blood-derived proteins and surface conditioning were not explicitly controlled and may influence crystal violet signals.

### ECC harbor iron acquisition and metabolism genes

Circular genome maps were generated for the contigs harboring the highest number of iron-related genes in each strain. *E. asburiae* had the highest number (17) of iron-related genes on contig_2 (Fig. 5 A) while *E. roggenkampii* had more iron-related genes on contig_14(14) (Fig. 5 B). Genome annotation revealed ECC strains possessed diverse iron acquisition and metabolism determinants, including ferrous and ferric iron transporters, heme/hemin uptake systems, and siderophore biosynthesis and transport pathways. The identified genes include *feoABC* (ferrous iron transport), *fepABCDG*, *entABCDEF* (enterobactin synthesis and transport), *iroDE* (salmochelin-related genes) and heme uptake components such as *chuA/hemR.* These iron-related genes were clustered on the same contig and co-localized with loci associated with horizontal gene transfer, as predicted by AlienHunter.

**Fig. 5.**
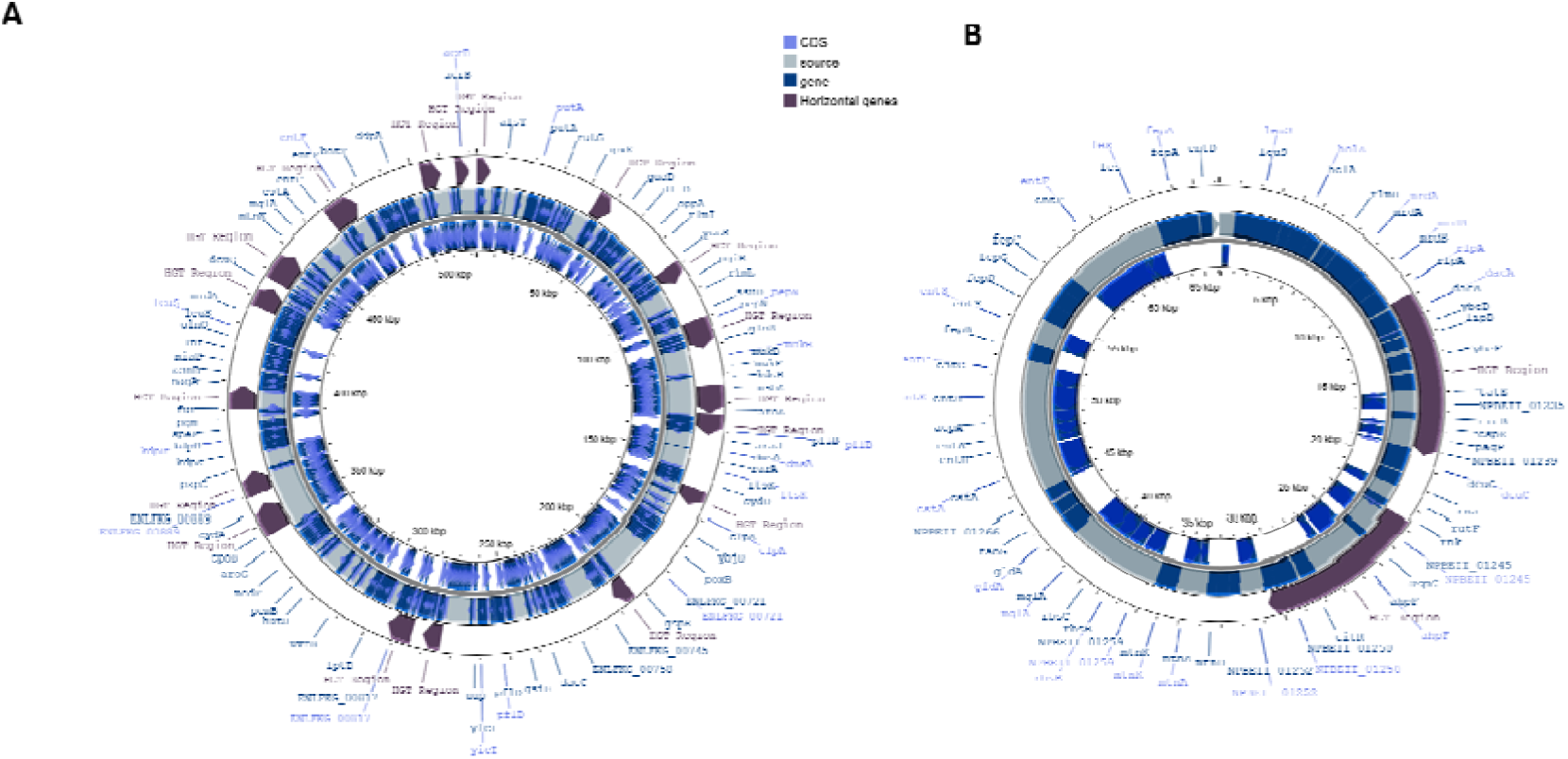
Circular visualization of iron rich contig in *E. asburiae* (A) and *E. roggenkampii* (B) using Proksee. *Coding sequences (CDSs) are shown as arrows indicating gene orientation. Annotated genes are highlighted in green, while the contig backbone is represented in pink. Dark purple regions indicate predicted horizontally transferred genomic regions (HGT). Inner rings represent GC content and GC skew*.

### ECC harbor resistance and virulence determinants

Genome analyses and annotation show *E. asburiae* encodes a higher number of enterobactin and salmochelin-associated genes as compared to *E. roggenkampii* which has increased aerobactin biosynthesis genes and prophage elements as shown in Fig. 6A. Comparative analysis of antimicrobial resistance genes revealed a shared core set of resistance regulators (*baeR, baeS, FosA, CRP, HNS*) present in both strains (Fig. 6B). The strains harbored resistance-associated genes including *qnrE* and *ramA*, suggesting distinct resistance determinants or global regulatory elements associated with baseline antimicrobial tolerance, while unique genes may reflect strain-specific adaptations. Genome annotation further revealed that both strains possess a broad repertoire of virulence-associated genes, including biofilm formation and motility-related genes. However, some of these functions were not consistently detected in VFDB BLAST searches, likely due to sequence divergence in draft genome assemblies rather than true absence.

**Fig. 6.**
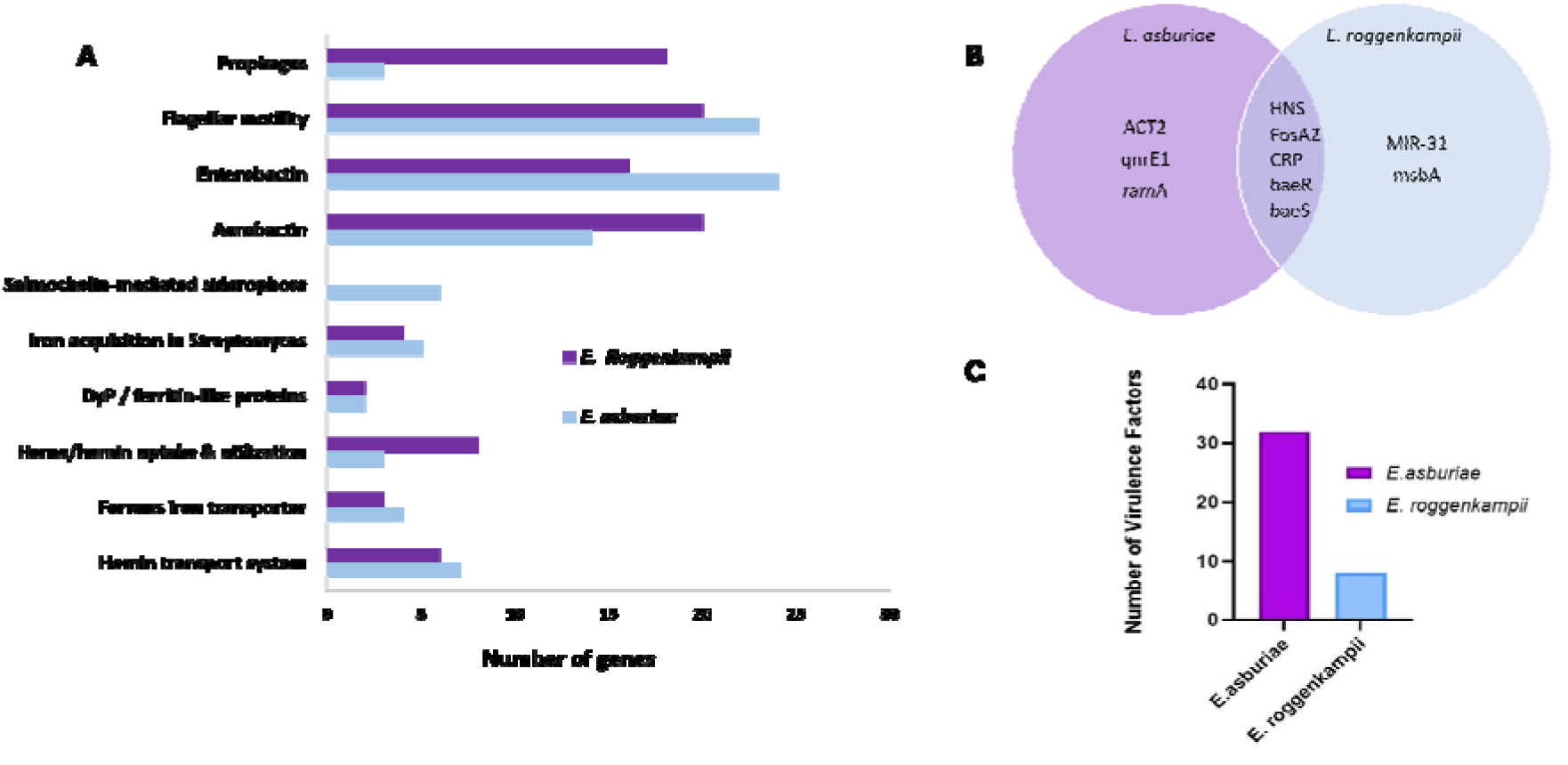
Comparative distribution of iron acquisition, siderophore biosynthesis, motility, and prophage-associated genes in *E. asburiae* and *E. roggenkampii* (A). Comparative analysis of resistance genes and virulence factors identified. ECC strains demonstrate shared and strain-specific antimicrobial resistance genes (B). *E. asburiae* harbors a greater number of VFDB-confirmed virulence factors than *E. roggenkampii* (C).

## Discussion

ECC strains exhibited enhanced growth with increasing iron concentrations in blood-rich media, suggesting the presence of iron acquisition systems that may facilitate utilization of host-derived iron during infection. This is consistent with studies linking bacterial iron utilization to pathogenicity (Zughaier & Cornelis, 2018). Intracellular iron quantification confirmed active uptake, with an initial increase followed by a decline under iron-replete conditions, while EDTA maintained low intracellular iron levels. This supports dynamic regulation of iron homeostasis during growth (Andrews et al., 2003). The observed decline may reflect repression of iron uptake, potentially mediated by the ferric uptake regulator (Fur), and a protective response against iron-induced oxidative stress (Carpenter & Payne, 2014). Ferric iron uptake systems have been implicated in the pathogenicity of *Pseudomonas aeruginosa* and may similarly contribute to virulence in the *Enterobacter cloacae* complex (ECC) (Pasqua et al., 2017).

Iron exposure markedly increased PMB MIC[[, with *E. roggenkampii* showing a 128-fold increase. This is consistent with reports linking altered iron homeostasis to antimicrobial resistance phenotypes (Choi et al., 2022; Méhi et al., 2014). These effects may reflect iron-mediated changes in bacterial physiology, including outer membrane composition and regulatory pathways implicated in polymyxin resistance in *Pseudomonas aeruginosa* and *Enterobacteriaceae* (Cheng et al., 2010), although such mechanisms were not examined here. A non-monotonic response to increasing PMB concentrations was also observed, similar to patterns reported in *Acinetobacter baumannii*, where higher antibiotic concentrations can select for tolerant subpopulations (Tsuji et al., 2016). The implications of this phenomenon remain unclear but may be relevant for treatment choices (Doijad et al., 2023; Fukuzawa et al., 2023; Tiseo et al., 2023). Iron availability was also associated with increased meropenem tolerance in a strain-dependent manner, consistent with reports of carbapenem tolerance *in E. cloacae*, although underlying mechanisms were not investigated (Murtha et al., 2022).

Iron-rich or iron-depleted conditions were associated with enhanced motility in ECC in a strain-dependent manner, although the increase was not statistically significant in *E. asburiae*. Iron limitation has been reported to increase bacterial motility, which is consistent with the enhanced motility observed under EDTA-mediated iron chelation (Burbank et al., 2015; Frick-Cheng et al., 2024; Nelson et al., 2019). Iron availability has also been linked to motility through siderophore-mediated iron acquisition systems (Burbank et al., 2015), which contribute to bacterial fitness and adaptation, although these mechanisms were not directly assessed in this study. In addition, previous studies have shown that iron levels do not necessarily repress siderophore-associated pathogenicity islands, such as those involved in yersiniabactin production (Magistro et al., 2017). The observed increase in motility under varying iron conditions may facilitate environmental adaptation and potentially contribute to the expression of virulence-associated traits in ECC. Similarly, iron availability influenced biofilm formation, with both iron-rich and iron-depleted conditions enhancing biomass in a strain-dependent manner. These findings are consistent with reports that iron availability and limitation can modulate biofilm formation across bacterial species (Chen et al., 2020; Oliveira et al., 2021; Wu & Outten, 2009).

Genomic features were consistent with observed phenotypes, with the presence of iron acquisition, heme utilization, and siderophore-associated genes aligning with enhanced growth under iron-rich conditions. Co-localization of iron-related and biofilm-associated genes suggests potential coordinated regulation of these functions. Additionally, the presence of MDR efflux systems and regulatory genes, including RND *(acrA/acrB)* and MFS transporters, may contribute to the observed antimicrobial resistance (Guérin et al., 2016). Differences in resistance and virulence-associated genes further support strain-specific pathogenic potential. Collectively, these findings highlight iron availability as a key environmental factor influencing multiple virulence-associated traits in ECC, including growth, antimicrobial tolerance, motility, and biofilm formation.

## Conclusion

Our findings show that iron supplementation is associated with increased growth, motility, biofilm formation, and reduced antibiotic susceptibility in *Enterobacter cloacae* complex. These phenotypic observations are supported by genomic analyses identifying multiple iron acquisition and metabolic genes. Collectively, the results suggest that iron metabolism plays a role in modulating virulence-associated and survival traits in ECC under varying iron conditions. While these findings highlight the potential for iron availability to influence pathogenic potential, further studies are required to elucidate the regulatory mechanisms linking iron sensing, gene expression, and adaptive responses during infection. In addition, the mechanisms of iron utilization were not directly assessed in this study; intracellular iron levels following red blood cell co-incubation were not quantified, and heme availability was not experimentally controlled, indicating that increased iron acquisition from erythrocyte lysis remains inferential.

## Acknowledgements

The authors appreciate AMR Research Group led by Dr. Abiola Isawumi at West African Centre for Cell Biology of Infectious Pathogens, Department of Biochemistry, Cell and Molecular Biology, University of Ghana for supports.

## Acknowledgement of Funding

This research was funded by the Gates Foundation (GF) (INV-050873-Amenga-Etego). The work was also supported by the National Institute of Health and Care Research (NIHR) [#134717] using UK international development funding from the UK Government to support global health research. The views expressed in this publication are those of the author(s) and not necessarily those of the GF, NIHR or the UK government. Also, a World Bank African Centres of Excellence grant (WACCBIP+NCDs: Awandare) and a DELTAS Africa grant (DEL-22-014: Awandare).

## Conflict of Interest

The authors report no conflict of interest

## Data Availability Statement

All the data generated have been included in the manuscript and the supplemental data.

## Notes

### Competing Interest Statement

The authors have declared no competing interest.

